# Perceived Timing of Postural Instability Onset With and Without Vision

**DOI:** 10.1101/2023.07.28.550867

**Authors:** Robert E McIlroy, Michael Barnett-Cowan

**Affiliations:** Department of Kinesiology and Health Sciences, University of Waterloo, 200 University Avenue West, Waterloo, Ontario, Canada, N2L 3G1

## Abstract

**Background:** Vision can significantly impact both the perception and behaviour related to postural control. This study examines the influence of vision on the perception of postural instability onset. Previous research employing Temporal Order Judgment (TOJ) tasks to investigate the perceived timing of postural perturbation onset has not incorporated visual cues.

**Research question:** Does the presence of visual feedback affect the point of subjective simultaneity (PSS) between postural perturbation onset and an auditory reference stimulus and does this additional sensory cue increase TOJ precision?

**Methods:** Using a lean-and-release paradigm, 10 participants were exposed to postural perturbations in both eyes closed (EC) and eyes open (EO) conditions using a TOJ task where they indicated whether they perceived postural instability onset or sound onset as occurring first for each trial. Separate paired t-tests between EC and EO PSS and just noticeable difference (JND) values were used. One-sample t-tests were also used on PSS values for both conditions, comparing them to 0ms (true simultaneity).

**Results:** The EC condition demonstrated a perceived delay of postural instability onset by 25.78 ms, while the EO condition showed a perceived delay of the auditory stimulus by 12.33 ms. However, no significant differences were found between the conditions or in comparison to true simultaneity. Mean JND values for EC (39.88 ms) and EO (46.48 ms) were not significantly different, suggesting visual information does not affect response precision for this task under these conditions.

**Significance:** These findings indicate that visual information does not significantly affect the perception of postural instability onset. This suggests that visual information may play a limited role in the early perceptual stages of postural instability.

## 1. Introduction

Perception of sensory stimuli significantly shapes our interaction with the world, including decisions and actions. The Central Nervous System (CNS) detects and processes sensory cues, which contribute to decisions like maintaining posture. This requires integration of proprioceptive, visual, and vestibular systems to model body motion relative to gravity. Speed discrepancies across sensory modalities introduce perceptual challenges, as the CNS must link related signals despite temporal asynchronies. In response to a threatening event such as a postural perturbation, being able to rapidly maintain balance is important for survival and how the event is perceived could help with perceptual learning to reduce falls.

Each sensory modality has different transduction times to generate a neural signal, which then travels at different speeds and distances before arriving at the cortex. Consequently, the CNS must determine which signals are related to the same sensory event despite temporal asynchronies. Research has shown that humans can perceive multisensory stimuli as simultaneous despite these differences in temporal processing across sensory modalities [1-3]. Our lab’s early studies demonstrated that when actively perturbing balance, the postural perturbation needs to significantly precede an auditory cue to be perceived as simultaneous when measured with a temporal order judgment (TOJ) task [4,5]. These findings suggested that the perceived onset of postural instability is slow. However, our most recent work suggests that these perceptual delays for fall onset depend on methodological design of the psychophysical experiment used to measure them [6]. After resolving the contribution of methodological design, we now consider the role of visual information in the perceived timing of postural perturbation as all past work has had the eyes closed.

Visual information is known to be processed more slowly compared to auditory and proprioceptive cues [7-9]. Visual information has also been observed to have perceptual delays compared to auditory stimuli [10-12]. During a temporal order judgment (TOJ) task between active head motion and an auditory cue, the presence of visual cues exhibits a tendency to reduce the point of subjective simultaneity (PSS) and just noticeable difference (JND) but does not statistically alter these measures with visual cues present [13]. Despite the abundance of sensory information, the presence of visual cues does not improve perception speed or precision of temporal order judgments. This insight contributes to the theory that visual information might not be critical in the early stages of balance responses. However, real-world scenarios often involve integrating all senses, potentially affecting postural control differently than in limited sensory situations.

In this study, we examined the effect of visual information on perceiving postural instability onset, using a TOJ task to determine the PSS and the JND [14]. The role of visual availability was assessed with two conditions, eyes open (EO) and eyes closed (EC) conditions.

Our hypotheses were: availability of visual information during postural perturbation would not significantly change the PSS [4-6]; onset of postural instability would not be perceived more slowly compared to an auditory stimulus in both EO and EC conditions [7,15,16]; and the JND would decrease in the EO conditions due to the observed effects of visual cues on postural sway and perceptual upright tasks [17-19].

## 2. Methods

### 2.1 Participants

A total of 13 participants were recruited for participation within the study. Two participants were excluded from analysis: one participant opted out before completion of all trials while the other did not follow response instructions correctly and treated the task as a reaction time test. Resulting in eleven healthy young adults (5 women; 21-26 years) data being utilized within this study. Participants reported no known musculoskeletal, auditory, visual, vestibular, or neurological disorders that may have affected their ability to perform the task. The study was conducted in accordance with the guidelines of the University of Waterloo Research Ethics Committee, and informed written consent was obtained from all participants after providing them with detailed information about the study procedures.

### 2.2 Protocol

The methodology utilized within this study was consistent with that of previous research conducted within our lab [6]. Participants made temporal order judgment (TOJ) responses to a postural perturbation evoked by a lean-and-release mechanism and an auditory reference cue. The lean-and-release and auditory reference stimulus setup was identical to that of previous research [6]. Participants continued to make their decisions of which stimuli occurred first using two handheld buttons. However, the two conditions utilized within this study design were eyes closed (EC) and eyes open (EO). During the EO trials, participants were instructed to look at an ‘x’ marked on a wall approximately 3 meters away [20]. During EC trials, participants wore a blindfold while also closing their eyes to ensure that they did not receive any visual cues if they were to accidentally open their eyes during the trial. Participants completed 144 trials (72 trials with vision and 72 trials without vision), with each SOA being repeated eight times. The SOAs were: −200 ms, −100 ms, −50 ms, −25 ms, 0 ms, 25 ms, 50 ms, 100 ms, 200 ms [6]. Negative SOAs indicate that the postural perturbation occurred first, and positive SOAs indicate that the auditory reference stimulus occurred first. SOAs were fully randomized for each participant, and after 24 consecutive trials, participants were given a 3-minute break to sit down to avoid fatigue over the course of the study. Each trial lasted for 15-20 seconds, with a comparable delay between each trial. Participants were told to relax while keeping their arms and hands comfortably at their sides throughout the trials. Participants were then taken through five practice trials.

Participants made TOJ responses to a postural perturbation evoked by a lean-and-release mechanism and an auditory reference stimulus produced through headphones. Participants were first weighed to determine the 7-8% body mass lean angle that was adopted throughout the study. Participants were then fitted with a full body harness that allowed for the attachment of the lean cable at the level of the 2nd and 3rd thoracic vertebrae and the safety rope, which was secured to the ceiling to prevent injury in case of an inability to recover balance [4,5,20,21]. The participants were positioned approximately 1 m from the lean-and-release apparatus, and their feet were positioned in a standardized position (heel centers 0.17 m apart, 14° between the long axes of feet [4,5,22]. The ground was marked with tape along the lateral borders of the feet, and a piece of wood was used as a heel stop to ensure the foot position was not altered during the study. Participants were not given any specific instructions on how to respond to the postural perturbation to prevent any voluntary changes that could occur to the postural response adopted by the participants. However, while the lean angle of 7-8% body mass produced a large enough perturbation to inherently evoke a stepping response in each of the subjects, it was not small enough to allow for a fixed support postural strategy [4,5,21,23,24].

### 2.3 Lean and Release Apparatus & Stimuli

As stated, the lean and release mechanism and the auditory stimuli were identical to that used in previous research; please refer to this research for specific characteristics of the system [6].

### 2.4 Data Analysis

Analysis of the TOJ responses was identical to that utilized in previous research [6], wherein the binary responses (0: postural perturbation occurred first; 1: auditory cue occurred first) were averaged at each of the specified SOAs. Negative SOAs represent that the postural perturbation occurred first, and positive SOAs represent that the auditory cue occurred first. A two-parameter logistic function (Eq. 1) was fitted to each of the participants’ averaged responses as a function of SOA using SigmaPlot 12.5.

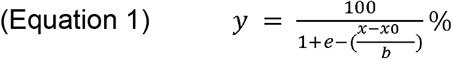

Where the inflection point of the logistic function (x0) was taken as the point of subjective simultaneity (PSS) and the standard deviation (b) was taken as the just noticeable difference (JND) [4].

Outcome parameters were inspected to identify potential outliers or poor-quality fits. If both the *R*^2^ of the logistic regression fit was ≤ 0.5 [25] and the p-value was *p* ≥ 0.05, then we would remove the participant’s data from the analysis, as we could not be confident that the logistic regression provided an accurate fit.

### 2.5 Statistical Analysis

We used a within-subjects’ design to assess the hypothesis that postural instability would not be perceived significantly slower than the auditory reference. A one-sample t-test was conducted comparing the mean PSS value of the EO and EC conditions to true simultaneity (0 ms). The one-sample t-tests specifically assessed whether the mean PSS values were significantly different from zero. To investigate the effect of visual feedback on mean PSS, a paired-sample t-test was performed to compare the EC and EO conditions. If normality failed in any comparisons as assessed by the Shapiro-Wilk test, an attempt was made to normalize the data. If these attempts with Z-Score normalization were not successful, then nonparametric statistics were performed. For each of the statistical tests, a significance level of α = 0.05 was utilized.

## 3. Results

One participant violating conditions for PSS and JND data was excluded due to their data fit falling outside *R*^2^ ≥ 0.5 and *p* ≤ 0.05 (EO: *R*^2^ = 0.377, *p* = 0.079; EC: *R*^2^ = 0.378, *p* = 0.078).

Figure 1 displays logistic fits for the remaining participants in both conditions. The mean PSS for EC (PSS mean = −25.78 ms, SE = 12.40) and EO (PSS mean = −12.33 ms, SE = 9.44) conditions were negative, suggesting postural instability needed to occur before the auditory cue to appear simultaneous (Figure 2). However, neither EC (*Z* = −1.478; *p* = 0.08) nor EO (*t*(9) = −1.307; *p* = 0.112) conditions demonstrated PSS values significantly less than zero, meaning visual absence or presence did not cause significant PSS change. The mean PSS difference between EC and EO was 13.44 ms, hinting that visual information slightly decreased PSS. Still, there was no significant difference between the mean PSS values of EC and EO (*t*(9) = 1.170; *p* = 0.272; 1 - β = 0.30), supporting our initial hypothesis that visual information would not change PSS significantly [13]. Notably, unlike previous research [6], EC did not yield a positive PSS mean.

**Figure 1:**
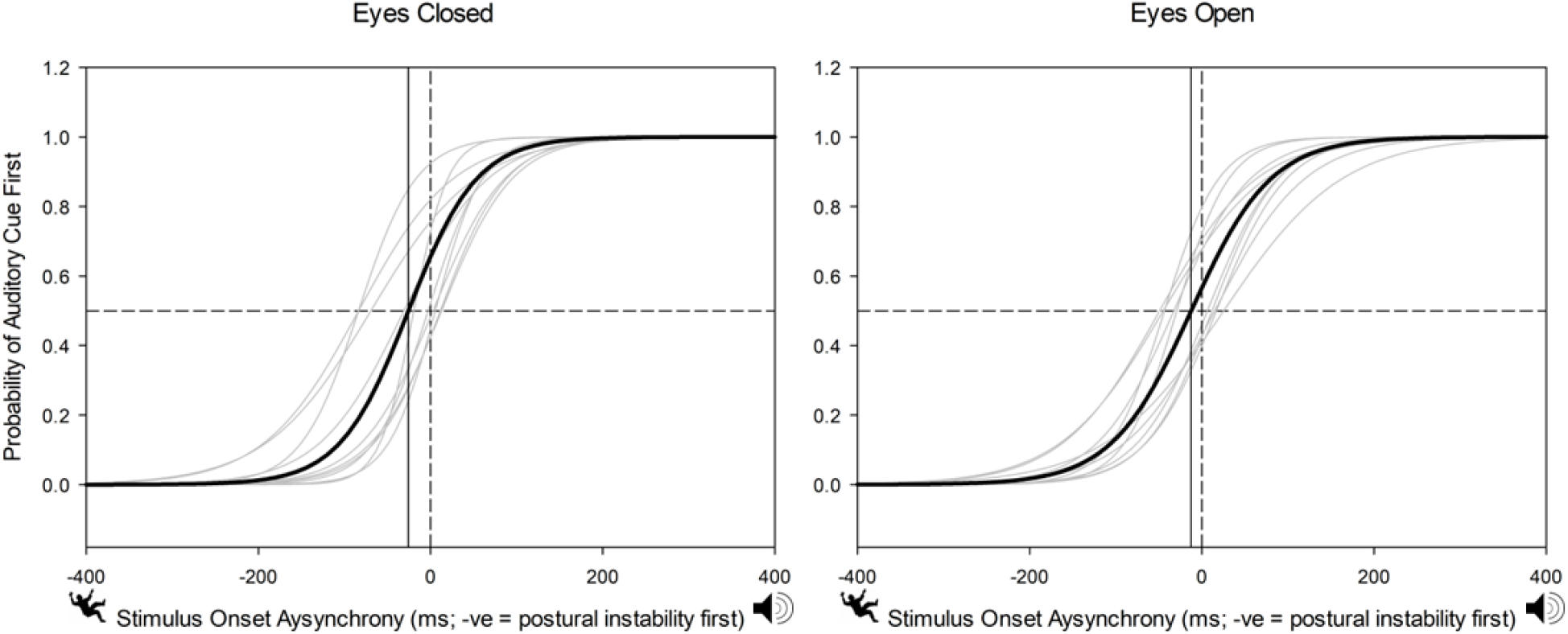
Logistic fits for both the Eyes Open versus Eyes Closed conditions. Light grey logistic function fitted lines represent each individual, while the black logistic function represents the mean of the individuals. The solid vertical line represents the mean PSS value for each condition. An SOA of 0 ms represents true simultaneity between stimuli and is exhibited by the vertical dashed line. The horizontal dotted line represents the 50% probability of the auditory reference stimulus occurring first, where the PSS is the time at which the logistic function crosses this 50% mark.

**Figure 2:**
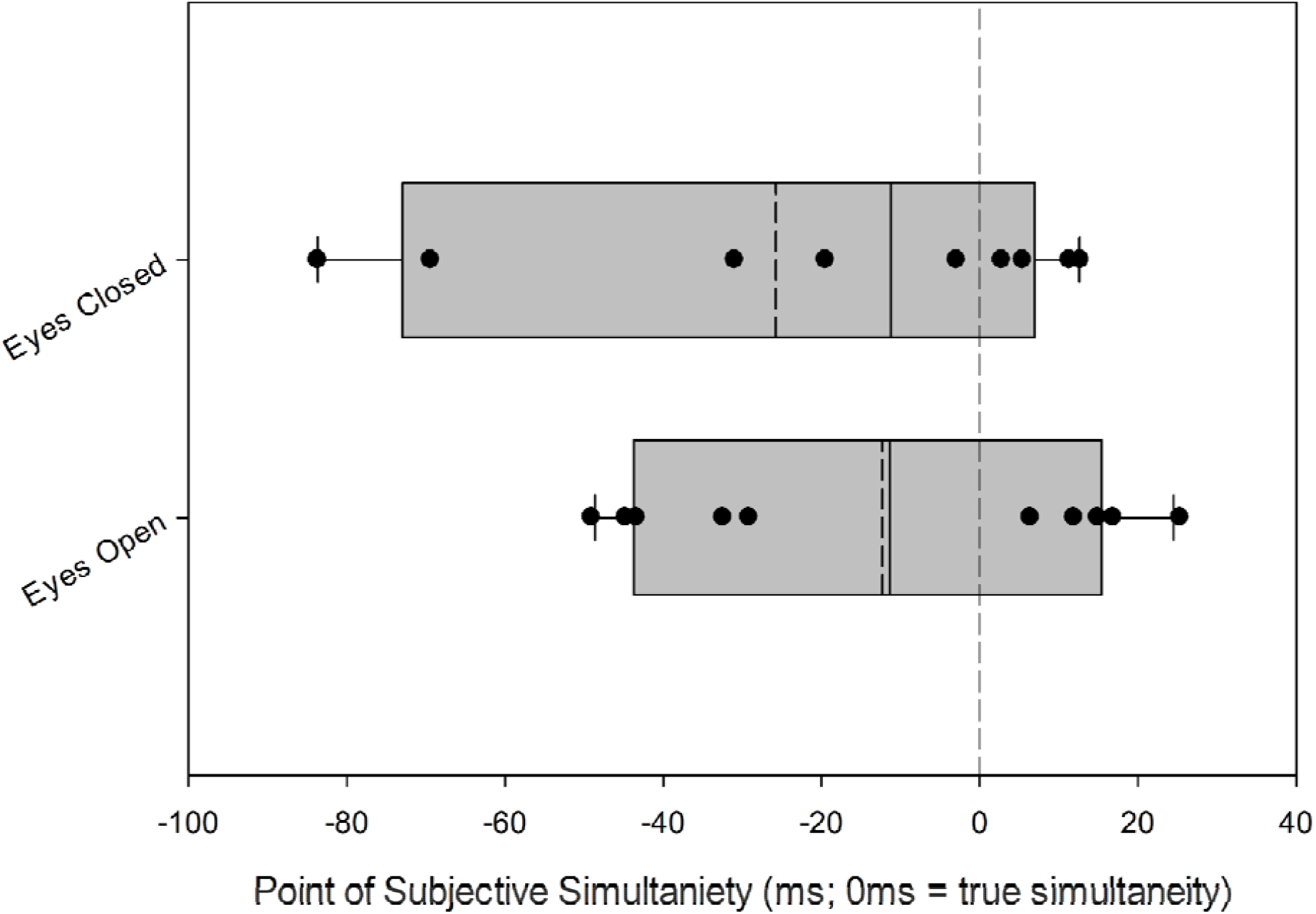
Mean PSS (dashed line), median PSS (solid line), and individual PSS values (circles) for the EC (top) and EO (bottom) conditions. Grey bars represent the 25-75th percentiles of the PSS data with error bars representing the 90th and 10th percentiles of data distributions. There were no statistically significant differences between the EC and EO conditions PSS values.

Mean JND values for EC (JND mean = 39.88 ms, SE = 38.04) and EO (JND mean = 46.48 ms, SE = 21.49) differed by 6.6 ms, a 16.5% change. Yet, a paired sample t-test revealed no significant differences (*t*(9)= 1.565; *p* = 0.152), suggesting visual information does not affect response precision. Standard error was larger for EC (Figure 3).

**Figure 3:**
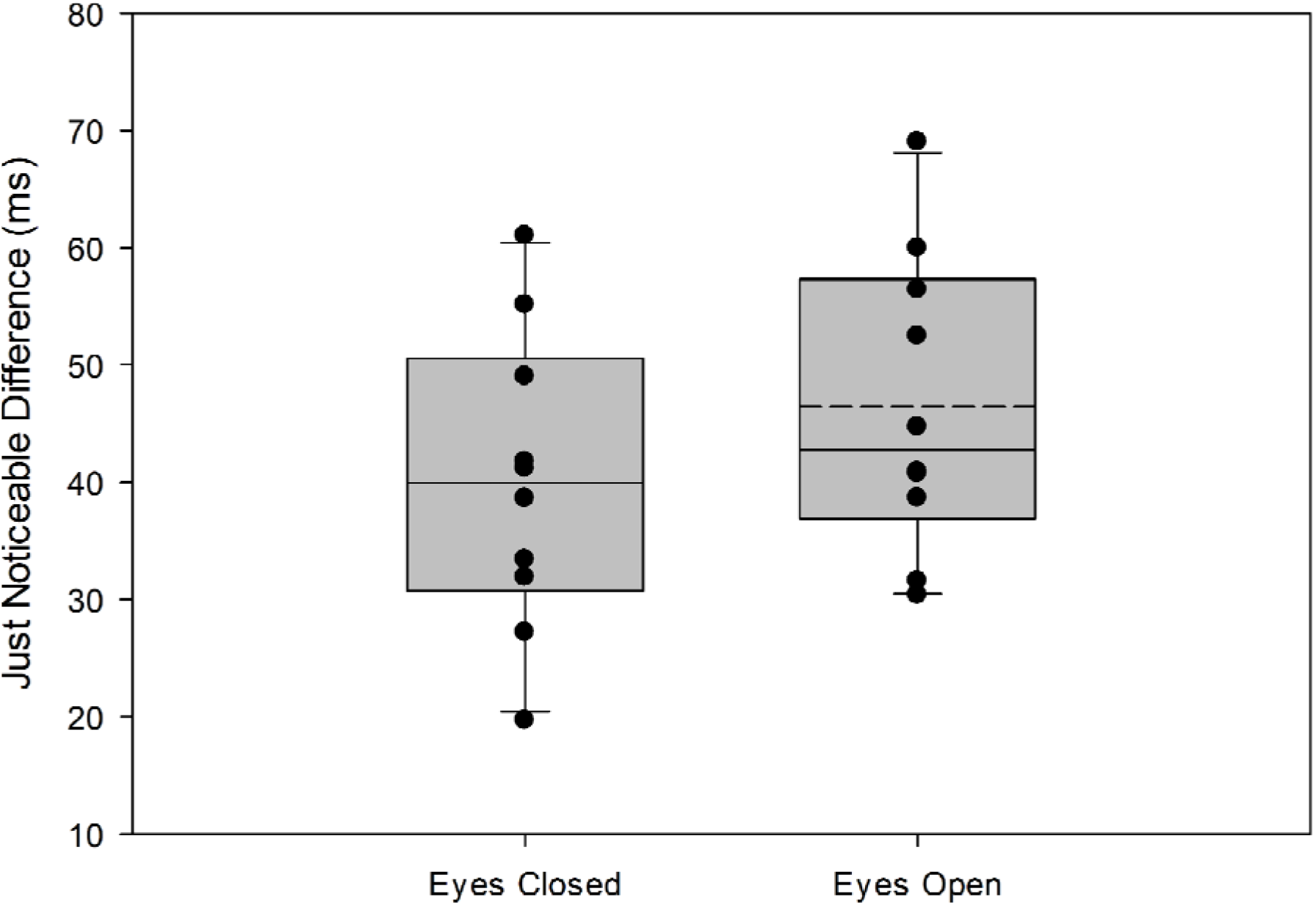
Mean JND (dashed line), median JND (solid line), and individual JND values (circles) for the EC (left) and EO (right) conditions. Grey bars represent the 25-75th percentiles of the JND data with error bars representing the 90th and 10th percentiles of data distributions. There were no statistically significant differences between the EC and EO conditions JND values.

Individual PSS and JND means varied significantly across participants for both conditions (Figures 1-3). Figure 1’s gray lines show logistic fits to each individual’s averaged responses at each SOA, depicting PSS and JND variability.

EC condition yielded a negative PSS mean (PSS mean = −25.78 ms, SE = 12.40), suggesting postural instability needed to precede the auditory cue by 25.78 ms to seem simultaneous. EO condition also had a negative PSS mean (PSS mean = −12.33 ms, SE = 9.44). Neither EC (*Z* = −1.478; *p* = 0.08) nor EO (*t*(9) = −1.307; *p* = 0.112) conditions showed PSS values significantly less than zero, indicating vision’s absence or presence did not significantly affect PSS. These findings support our initial hypotheses of no significant difference from true simultaneity in both conditions.

## 4. Discussion

This study examined the impact of vision on perceiving simultaneity between postural instability onset and an auditory cue. Negative PSS averages in both the EC and EO conditions indicate postural instability should precede the auditory cue for perceived simultaneity, consistent with prior work [4-6]. Absence of significant PSS differences between conditions supports the hypothesis that vision does not significantly alter simultaneity perception [13].

Previous work demonstrated postural perturbations must precede an auditory stimulus by about 45 ms for perceived simultaneity [4,5], with equal SOA distribution recently showing no significant delay [6]. This study, using an identical equal SOA distribution and methodology, found no statistically significant PSS difference from true simultaneity, both with or without vision. These reproducible results reinforce the use of equal SOA distribution in preventing lag adaptation from asynchronous stimuli presentation [1,6,26-29]. While additionally exhibiting, the presence of visual information does not have a significant impact on the perception of instability onset.

Although visual information aids balance control, stability can be maintained without visual feedback, albeit with increased postural sway [30-32]. This highlights the importance of other sensory information, specifically proprioceptive feedback in the detection of postural instability and the control of balance. Proprioceptive feedback’s importance in detection of body orientation and position in combination with slower processing speeds of visual information might explain the absence of a significant effect on onset of perception between the EC and EO conditions.

No significant PSS and JND shifts between conditions suggest minimal visual input reliance during this task. Participant variability might be due to sensory re-weighting, previous experience, fear of falling, attention, or fatigue. Future studies should increase sample sizes and use qualitative measures to account for this variability. A post-hoc power calculation of 1 - β = 0.30 for the PSS comparison between conditions indicates a probable type II error. An a priori power calculation suggests a sample of 77 for a power of 1 - β = 0.95, indicating sample size as a limiting factor within this study.

Multisensory integration and sensory re-weighting, occurs in situations where the CNS adjusts sensory information utilization under circumstances of sensory alterations, such as when vision is absent [33]. In the absence of visual information, the CNS will naturally adapt to upregulate the reliance on other sensory modalities such as proprioceptive cues. Proprioceptive information is heavily relied upon for posture and balance [33,34] and even more so in situations of lack of visual information. In circumstances of visual cue availability, it is a possibility that visual cues may not have been upregulated due to their relevance. Evocative visual scenes can impact postural responses [35-38], re-emphasizing that visual feedback and the visual scene presented to individuals is important in the control of posture.

A lean-and-release paradigm has limitations, potentially introducing learning effects or central set changes, due to the predictability of the perturbation direction. Future studies should use randomized perturbation strategies, like moving platforms, to limit these effects and explore different sensory inputs in postural instability perception.

In summary, vision does not significantly influence postural instability and auditory cue simultaneity perception. These findings enhance multisensory integration literature and illuminate vision’s role in auditory and postural information processing. Future work should investigate attention, cognitive load, and individual differences impacts on simultaneity perception, along with proprioceptive feedback’s role in postural instability onset perception using sway-referenced platforms or non-compliant surfaces. Additionally, exploring how visual context may alter onset perception could establish how visual information may be utilized in postural and perceptual tasks.

## Acknowledgements

Supported by a Natural Sciences and Engineering Research Council of Canada (NSERC) Discovery Grant (RGPIN-03977-2020) and an Ontario Early Researcher Award to MB-C.

